# Bacterial wilt resistance is correlated with rhizosphere bacterial communities in wild potato *Solanum malmeanum*

**DOI:** 10.64898/2026.02.11.705323

**Authors:** María Virginia Ferreira, Florencia Tourné, Ignacio Eastman, María Cecilia Rodríguez-Esperón, Gustavo Rodríguez, Matías González-Arcos, Francisco Vilaró, Guillermo Galván, Paola Gaiero, Giovanni Larama, Máximo González, Raúl Platero, María Inés Siri

## Abstract

Wild potatoes are long-evolving relatives of the cultivated potatoes we have today. These wild *Solanum* species harbor traits that can be exploited to develop more nutritive and resilient potato varieties, providing the genetic basis for resistance to abiotic and biotic stresses such as drought, low temperatures, diseases and pests. Wild potato species are widely used as valuable genetic resources in breeding programs, including efforts aimed at improving resistance to bacterial wilt caused by *Ralstonia solanacearum*. Among the wild species native to Uruguay, *Solanum malmeanum* has emerged as a particularly valuable source of resistance. The aim of this work was to investigate weather differences in bacterial wilt resistance among *S. malmeanum* accessions are associated with structural and compositional changes of rhizosphere bacterial communities. Two *S. malmeanum* accessions were compared, one susceptible (RN9P2) and the other resistant (A11P1) to bacterial wilt. The impact of plant resistance and pathogen colonization on the structure of rhizosphere bacterial communities were evaluated using high throughput 16S rRNA gene amplicon-sequencing. Significant differences were observed between accessions and pronounced shifts in rhizosphere bacterial communities were detected in response to pathogen inoculation. *Cryseobacterium*, *Sphingobacterium, Komagataeibacter, Gluconobacter, Lactobacillus* and *Dyella* were differential genera and enriched in the rhizosphere of the resistant accession. Several of these genera have been previously associated with disease suppression. Overall, these results suggest that the rhizosphere bacterial community associated with resistant *S. malmeanum* accessions may contribute to protection against *R. solanacearum* infection.

## 1. Introduction

Potato (*Solanum tuberosum* L.) is classified as the fourth most important cultivated food after rice, corn, and wheat and is consumed by over a billion people worldwide (Devaux et al., 2014). It belongs to the Solanaceae family and *Solanum* genus, which comprises about 2000 species, while section Petota (where potato and its wild relatives belong) contains 112 species (Spooner et al., 2014; Gagnon et al., 2022; Bozan et al., 2023). South America is the main center of origin for many potato species. Even though cultivated potatoes have high sequence diversity, early improvement was based on a limited gene set (Hardigan et al., 2017), therefore it is necessary to study landraces and related wild species to introduce new agricultural traits of interest. Currently, there are 107 recognized wild potato species (Gagnon et al., 2022), representing an enormous source of untapped adaptive potential that can be used to broaden the genetic basis of the crop (Hardigan et al., 2017), creating more nutritious potato varieties and introgressing genes for resistance to different stress factors such as drought, low temperatures, diseases, and pests (Spooner and Hijmans, 2001; Loginov et al., 2018). Potatoes are susceptible to a series of diseases, the most important are late blight caused by *Phytophthora infestans* L., bacterial wilt caused by *R. solanacearum*, and various viral diseases (Weng et al., 2024).

*R. solanacearum* is the causal agent of bacterial wilt, considered one of the most economically impactful plant diseases worldwide and second most important bacterial plant pathogen (Elphinstone, 2005; Mansfield et al., 2012). This pathogen affects more than 250 plant species within 50 families. In addition to its broad host range, *R. solanacearum* has a wide geographic distribution, affecting species in tropical, subtropical, and temperate zones (Genin and Denny, 2012; Peeters et al., 2013; García et al., 2019). Uruguay has a temperate climate, with potato crops being highly affected by *R. solanacearum* (Pianzzola, 2012). In colder climates, strains of *R. solanacearum* are characterized by high virulence and the ability to produce asymptomatic latent infections (Cellier and Prior, 2010).

Currently, there is no effective control method against *R. solanacearum.* The best strategy involves integrated management by combining different components or control strategies. These methods include the use of resistant or tolerant varieties, the application of good agronomic practices, as well as the adoption of biological and chemical control strategies. Additionally, using certified pathogen-free seed with a crop rotation scheme also contributes to limiting the presence of the pathogen (Yuliar et al., 2015). The introduction of resistance to *R. solanacearum* in the host plant is a crucial component for achieving long-term disease control, as it is the most sustainable, cost-effective, and environmentally friendly strategy (Boshou, 2005). Some wild potato species have been described as highly tolerant and, therefore, considered potential sources of resistance in breeding programs. A number of wild potatoes species with some level of resistance to bacterial wilt have been reported, such as: *S. acaule, S. andreanum, S. brevicaule (as S. brevicaule s.s. and S. sparsipilum), S. bulbocastanum, S. candolleanum, S. cardiophyllum, S. chacoense, S. clarum, S. commersonii, S. jamesii, S. malmeanum, S. microdontum, S. pinnatisectum, S. tuberosum* Group Phureja*, S. tuberosum* Group Stenotomum (Boshou, 2005; González et al. 2013; reviewed by Machida-Hirano 2015; Andino et al. 2022; Norman et al. 2020; Nicolao et al. 2022; Núñez et al. 2023; Stancov et al. 2023, reviewed by Sarkinen et al. 2025). The most valuable wild species available in Uruguay are *S. commersonii* and *S. malmeanum.* These tuberous species are widely distributed in the country and exhibit significant morphological, chemical, and genetic diversity, as well as important levels of resistance to *R. solanacearum* (Vázquez et al., 1997; Siri et al., 2004, 2011; Ferreira et al., 2017). For several years, the national potato breeding program has been working on the introduction of resistance to bacterial wilt from *S. commersonii* and *S. malmeanum* into new potato varieties adapted to local environmental conditions. Genotypes from the breeding program with differential responses to *R. solanacearum* infection were selected and characterized (González et al., 2013; Ferreira et al., 2017; Gaiero et al., 2017; Andino et al., 2022; Ferreira et al., 2024).

In addition to plant genetic resistance, increasing evidence indicates that soil microorganisms and plant-associated microbiota play a key role in plant health and disease resistance. The rhizosphere is the narrow zone of soil surrounding and influenced by plant roots. The rhizosphere microbiome is defined as the community of microorganisms present in the rhizosphere, including the molecules and metabolites they produce, structural elements and interactions with the environment (Mendes et al., 2013; Berg et al., 2020). Microorganisms that colonize the rhizosphere can provide beneficial functions for the host plant, particularly in nutrient acquisition, stress tolerance, and protection against soil-borne pathogens (Mendes et al., 2013; Pérez-Jaramillo et al., 2016). Rhizosphere microbiome diversity depends on the nutrients released by host plants, which can be influenced by biotic and abiotic factors, including environmental properties (soil type, climatic conditions, and agricultural practices), the plant developmental stage, and the plant species itself (Nannipieri et al., 2003; Berg and Smalla, 2009; İnceoğlu et al., 2011; Mardanova et al., 2019). Additionally, root exudates may be related with plant genotypes, leading to differential bacterial communities associated with the rhizosphere among genotypes of the same species (Berg and Smalla, 2009; İnceoğlu et al., 2011). Proteobacteria and Actinobacteria are reported as the most abundant phyla of the potato rhizosphere (Roquigny et al., 2018). However, significant differences in distribution and abundance of phyla have been observed in different cultivars, plant growth stages, and soil types on the potato rhizosphere microbiome (Milling et al., 2005; Van Overbeek and Van Elsas, 2008; İnceoğlu et al., 2010, 2011; Weinert et al., 2011; Barnett et al., 2015; Pfeiffer et al., 2017; Mardanova et al., 2019; Pantigoso et al., 2020).

The domestication process of cultivated crops, modified plant traits involved in recruiting and structuring their associated microbial communities. Such changes, together with the transition from natural habitats to agricultural soils, may have reduced the ability of cultivated varieties to assemble microbial taxa that support ecological functions relevant to plant health (González et. al., 2025). Development of plant diseases caused by soilborne pathogens is associated with plant defense responses, environmental conditions, and interactions with the rhizosphere microbiome associated with the host plant. Most reports of rhizosphere microbiome in plants infected by *R. solanacearum* have been conducted in tomato and tobacco. It has been demonstrated that pathogen invasion alters bacterial communities in the rhizosphere, leading to a clear reduction in diversity and abundance of beneficial microorganisms (Gu et al., 2016; Wang et al., 2017; Yang et al., 2017; Wei et al., 2018; Mao et al., 2020; Wen et al., 2020). Building on this evidence, we hypothesized that resistance of the wild potato *S. malmeanum* to *R. solanacearum* is influenced by the composition and functional potential of its rhizosphere bacterial microbiota, which may contribute to limiting pathogen establishment or disease progression. In this study, we compared the rhizosphere bacterial communities associated with resistant and susceptible *S. malmeanum* accessions under pathogen-inoculated and non-inoculated conditions, aiming to identify microbial taxa and functional traits potentially associated with resistance to *R. solanacearum*.

## 2. Materials and Methods

### 2.1. Bacterial strains and growth conditions

*R. solanacearum* strain UY031 (Siri et al., 2011; Guarischi-Sousa et al., 2016) was streaked from a glycerol stock at -80 °C on triphenyltetrazolium chloride (TZC) agar plates (Kelman, 1954), and incubated at 28°C for 48–72h. Liquid cultures were conducted in Kelman medium (1954) without triphenyltetrazolium. Optical density was measured spectrophotometrically at 600 nm to adjust bacterial suspensions for inoculation (OD_600_ of 0.1 corresponds to 10^8^ cfu/mL).

### 2.2. Plant materials and growth conditions

Two wild potato accessions of *S. malmeanum* with contrasting levels of resistance to bacterial wilt, RN9P2 (susceptible) and A11P1 (resistant) (Núñez et al., 2023; Stancov et al., 2023), were selected from a wild potatoes germplasm collection assembled in various field collection efforts led by our group at the College of Agriculture (Universidad de la República) and maintained at the National Research Institute for Agriculture (INIA) in vitro germplasm bank.

Plants were micro-propagated *in vitro* from a node in Murashige and Skoog medium supplemented with sucrose 30 g/L and maintained at 22°C with cycles of 16-h light/8-h darkness for 6 weeks in growth chambers. Subsequently, they were propagated by cuttings via a semi-autotrophic hydroponic (SAH) method (modified from Rigato et al., 2001), in plastic boxes filled with soil mix (Tref Substrates BV, Moerdijk, Netherlands) and kept at 22°C with cycles of 16-h light/8-hour darkness, for 2 weeks in growth chambers. Then, they were transferred to seedling trays, with one plant per cell, and maintained at 22°C with cycles of 16-h light/8-h darkness for 3 weeks in growth chambers. Plants were transplanted into 5 L pots with soil from a potato field with no previous history of bacterial wilt and were maintained for 4 weeks (until inoculation) in a high tunnel to achieve plant growth under natural field conditions, with natural light and temperature.

### 2.3. Plant inoculation assays

Bacterial suspensions of *R. solanacearum* UY031 (Siri et al., 2011) were prepared from overnight liquid cultures and spectrophotometrically adjusted to a concentration of 10^8^ cfu/mL of saline solution (OD600 of 0.1). Petri dishes (9 cm in diameter) containing TZC agar (Kelman, 1954) were inoculated with 100 μL of the prepared suspension, and confluent growth across the entire dish was achieved through the spreading plate technique. After 48 h of incubation, a piece (1/8) of the grown agar plate was buried in the soil at 20 cm depth and 5 cm away from the base of each plant (Fort et al., 2020). Disease progression was recorded until 27 days post inoculation (dpi) using an ordinal scale ranging from 0 (asymptomatic plant) to 4 (all leaves wilted) (Winstead, 1952). The resistance level was calculated by the area under disease progress curve (AUDPC) based on the average wilt scoring for each genotype. Analysis of variance (ANOVA) and the unpaired Student’s *t* test were applied with a 95% confidence level to analyze AUDPC. Model residuals were used to check for the assumptions of normality and homogeneity of variances. Statistical analyses were done using R software (Version 4.3.0). Non-inoculated plants were included in the assay as controls. Experiments were performed using 3 plant replicates for each genotype (susceptible and resistant) per treatment and sampling time.

### 2.4. Rhizosphere sampling and sample processing

Rhizosphere samples were collected at 13 and 27 dpi to analyze the microbiome composition at different disease stages. Samples of non-inoculated plants maintained under the same conditions were collected to evaluate the microbiome dynamics in healthy plants without pathogen contact.

For sample collection, plants were removed from the pots and non-adhered soil of roots was carefully discarded. The entire root system with the tightly adhered soil was weighed and placed in sterile stomacher bags (Seward Stomacher®, United Kingdom) with 100 ml of saline solution and treated with a stomacher (Seward Stomacher® 400 Circulator Lab Blender, United Kingdom) for 10 minutes at medium speed. Supernatant was collected after 2 minutes of centrifugation at 12,000 rpm for pathogen quantification and to harvest the rhizosphere. Rhizosphere pellet was stored at -80 °C until DNA extraction (Mardanova et al., 2019; Elsayed et al., 2020). Total environmental DNA (eDNA) was extracted from the rhizosphere pellet (400 mg) with the FastDNA Spin Kit for Soil (MP Biomedicals, Heidelberg, Germany). Purity and concentration of DNA samples were measured by Nanodrop (Genova nano, Jenway, United Kingdom) prior to storing at -20 °C for further analysis.

### 2.5. *Ralstonia solanacearum* rhizosphere quantification

Plate counting was used to quantify *R. solanacearum* concentration in the rhizosphere of inoculated plants. For this purpose, 1 mL of supernatant was taken before centrifugation, and serial dilutions were prepared. Colony counts were performed using semi-selective medium (mSMSA) plates (Elphinstone et al., 1996) incubated at 28°C for 2-5 days. Plate counts were represented as mean ± standard error of mean (*SEM*). Statistical analysis was carried out with unpaired Student’s *t* test and one-way analysis of variance (ANOVA). Model residuals were used to check for the assumptions of normality and homogeneity of variances. Statistical analyses were done using R software (Version 4.3.0).

### 2.6. Sequencing and analysis of 16S rRNA gene amplicons from total community DNA

Amplification, library construction, and sequencing were carried out by Novogene Corporation Inc. (USA). The set of primers used for amplification (341F/806R) flank the V3-V4 variable region of approximately 470 bp of the 16S rRNA gene of Prokaryotes including domains of Bacteria and Archaea (Klindworth, et al. 2013). Libraries containing 16S rRNA genes were analyzed using 2 × 250 bp paired-end high-throughput sequencing on an Illumina NovaSeq 6000 platform.

Reads obtained from sequencing of the V3-V4 variable region of 16S rRNA gene were analyzed using the QIIME2 software (Bolyen et al., 2019). Analysis involved quality control steps, forward and reverse sequence assembly, generation of representative sequences, amplicon sequence variants (ASV), using the DADA2 algorithm (Callahan et al., 2016), and removal of chimeric sequences. Taxonomic assignment for ASVs was performed using the Naïve Bayes classifier, implemented in the scikit-learn library of Python, and trained with the SILVA v138 database (Quast et al., 2013). Before diversity calculations, contaminant sequences associated with mitochondria and chloroplasts were removed.

For diversity analysis, ASVs were aligned using the MAFFT algorithm, and a tree was constructed with a root based on distances calculated from the alignment using the FASTTREE algorithm (Price et al., 2009). Prior to calculating alpha diversity indices, abundances were normalized by rarefaction. Alpha diversity was assessed by calculation of the Shannon index, Pielou’s index, the number of ASVs and the phylogenetic diversity index of the ASVs. Significance of each index was assessed using the Kruskal-Wallis test with a confidence level of 95%. Beta diversity was estimated through Principal Component Analysis (PCA), where abundances were normalized using the Aitchison distance. Significance of each factor and its interaction was evaluated using the PERMANOVA test implemented in the Adonis function in the vegan package, considering a *p* < 0.05.

UpSet plots were generated using the UpSetR package (v1.4.0) in R (v4.3.0) (Conway et al., 2017) to visualize shared and unique Amplicon Sequence Variants (ASVs) among resistant and susceptible genotypes under both control and inoculated conditions. An ASV was considered present in a given group if it was detected in at least two out of three biological replicates.

Differential genera were identified using a Zero-Inflated Gaussian Mixture Model (ZIGMM) in the *metagenomeSeq* package (v1.36.0) (Paulson et al., 2013), applying a >0.5% relative abundance threshold and a significance level of *p* < 0.05 after FDR correction.

Co-occurrence networks for both resistant and susceptible genotypes were constructed using the NetCoMi (v1.1.0) (Peschel et al., 2021) R package. Microbial interactions were inferred using the SparCC correlation method (correlation threshold = 0.6, *p* < 0.05) (Friedman and Alm, 2012) with low-abundance taxa (<0.001% relative abundance) filtered to reduce noise. Network modules and node-level metrics, including degree, closeness, betweenness, and eigenvector centrality, were calculated using the cluster_fast_greedy algorithm in igraph (v1.3.5) (Csardi and Nepusz, 2006).

Functional profiles were inferred using the FAPROTAX database (v1.2.4) (Louca et al., 2016), which predicts microbial metabolic and ecological roles from taxonomic data. ASV annotations from the SILVA database (v138) were mapped to functional groups.

## 3. Results

### 3.1. Disease progression and *Ralstonia solanacearum* colonization in *Solanum malmeanum* accessions

Bacterial wilt progression was monitored in *S. malmeanum* plants inoculated with *R. solanacearum* over a 27-day period (Fig. 1a). Disease progression was significantly different among genotypes as indicated by AUDPC analysis (*p* = 0.039). While the resistant genotype remained asymptomatic throughout the experiment, the susceptible genotype exhibited severe wilt symptoms, confirming the contrasting resistance levels of the two accessions.

**Fig 1.**
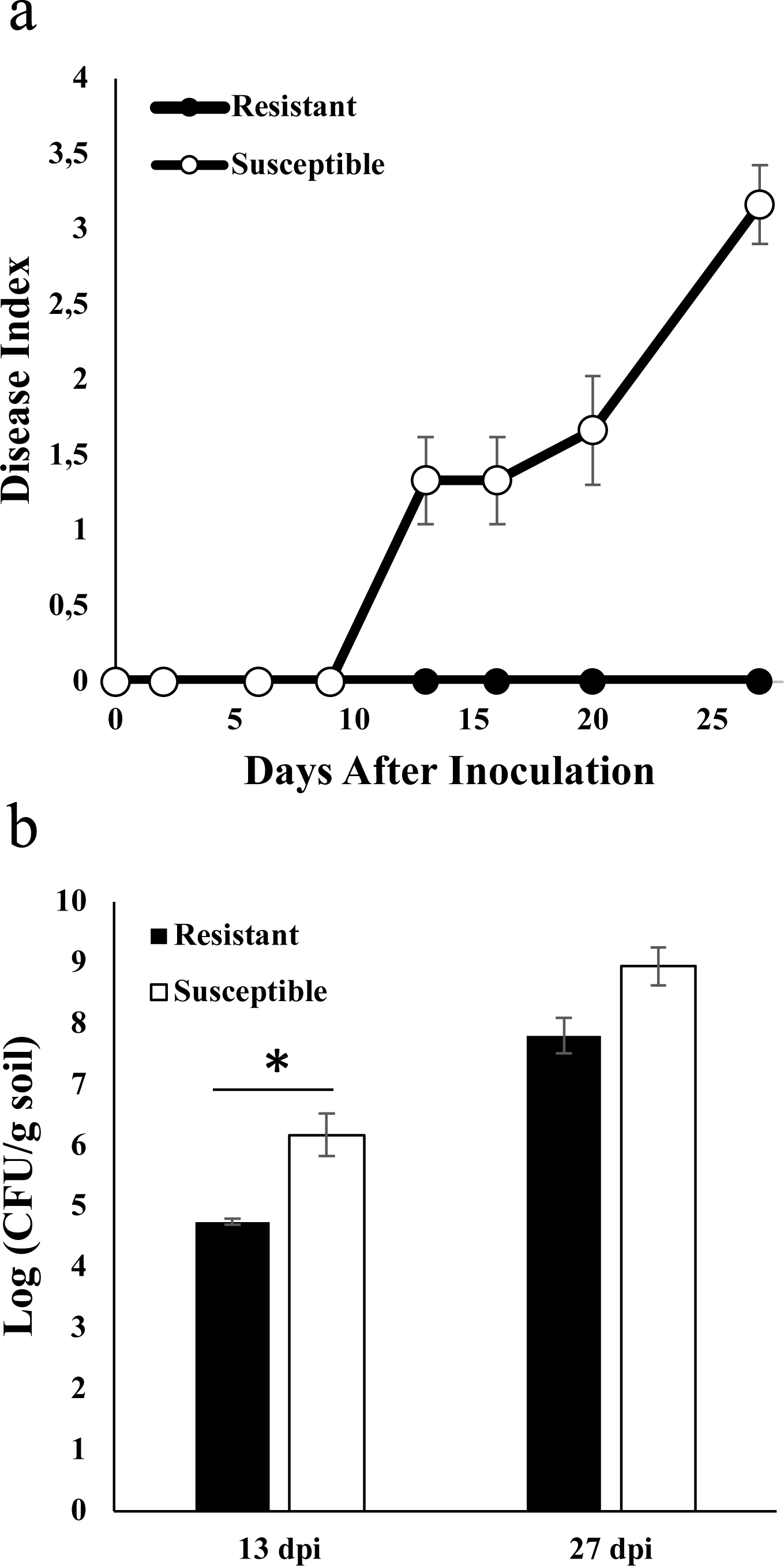
Bacterial wilt disease (caused by *Ralstonia solanacearum*) progression in the resistant and susceptible genotypes. Each data point represents the mean of three replicates. b. Population dynamics of *Ralstonia solanacearum* in the rhizosphere of the resistant and susceptible genotypes. Each bar represents pathogen concentration means (n = 3). * Indicates significantly different concentrations according to ANOVA and Student’s comparison test (p < 0.05). Vertical bars represent standard errors of the means.

*R. solanacearum* colonization was quantified in rhizosphere samples of inoculated plants at 13 and 27 dpi (Fig. 1b). The resistant genotype had a significantly lower concentration of pathogen (*p* = 0.015) than susceptible genotype at 13 dpi. At 27 dpi, pathogen populations in the resistant genotype increased to levels comparable to those observed in the susceptible genotype.

### 3.2. Effect of genotype and *Ralstonia solanacearum* infection on bacterial community structure in *Solanum malmeanum* rhizosphere

The effects of plant genotype and *R. solanacearum* infection on rhizosphere bacterial community structure were investigated in *S. malmeanum.* At 13 dpi, no significant differences in alpha diversity were detected between genotypes or treatments (Fig. 2a). At 27 dpi, alpha diversity showed increased variability among genotypes and treatments, however, no statistically significant differences were found (Fig. S1a).

**Fig 2.**
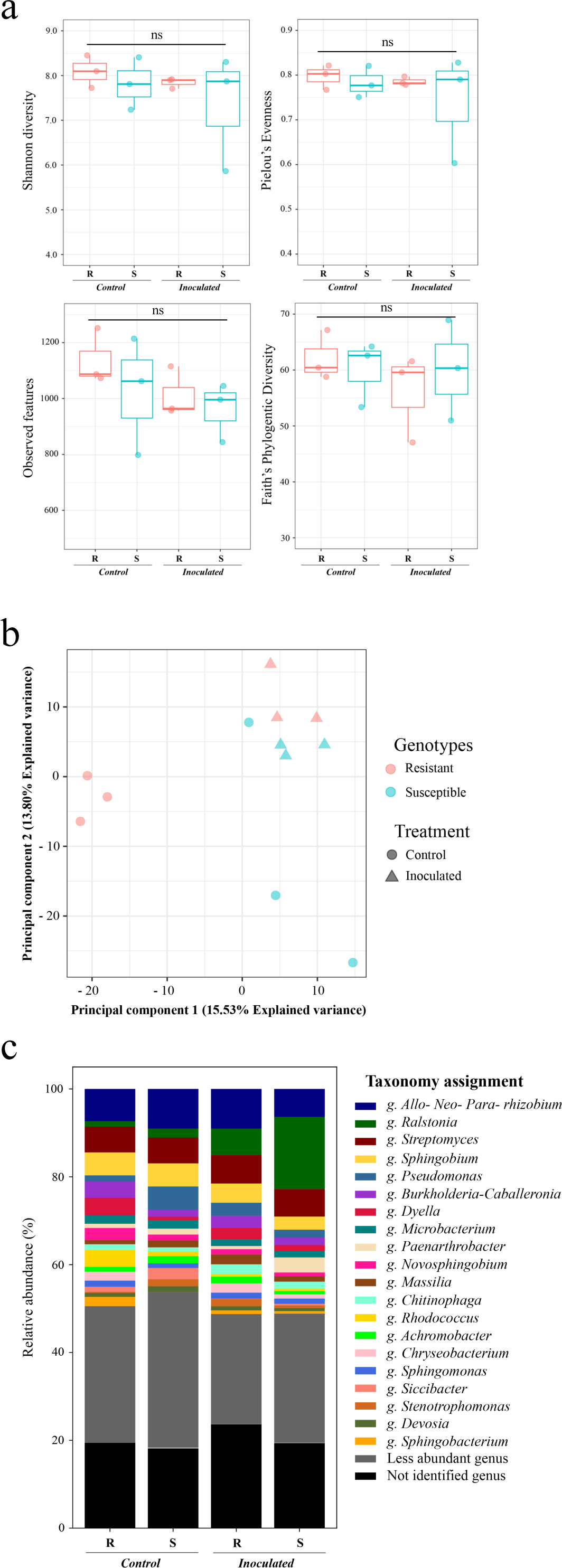
a. Comparative alpha-diversity analysis of rhizosphere bacterial community among the bacterial wilt resistant (R) and susceptible (S) genotype 13 days post inoculation. Significance of Shannon diversity index, Faith’s phylogenetic index, Pielou’s Evenness index and observed features (number of ASVs) were calculated with Kruskal-Wallis test (p < 0.05). ns = not significant. b. Comparative beta-diversity analysis of rhizosphere bacterial community between the bacterial wilt resistant and the susceptible genotype 13 days post inoculation. The distribution pattern of control and inoculated plants from the resistant and susceptible genotype were normalized using Aitchison distance. Significance of each factor and its interaction was evaluated using the PERMANOVA test (p < 0.05). c. Relative abundance (%) of the major bacterial genera in the rhizosphere microbiota of the resistant (R) and susceptible (S) genotypes 13 days post inoculation.

Beta diversity analysis revealed clear differences in rhizosphere bacterial community composition associated with both genotype and *R. solanacearum* infection. Principal Component analysis (PCA) using the Aitchison distance showed that the first two components explained 15.53% and 13.80% of the variability in the ASV log_10_ proportions, respectively (Fig, 2b). At 13 dpi, both inoculation (R^2^=0.1343; *p* = 0,001) and genotype (R^2^=0.1088; *p* = 0,001) had a significant effect on community composition. Additionally, the rhizosphere bacterial microbiota of plants at 27 dpi were distinct between genotypes (R^2^=0.1434; *p* = 0.011), but no significant differences were found between control and inoculated plants (Fig. S1b). Based on the results described above, subsequent analyses in this study focused on samples collected at 13 dpi.

Relative abundance of the 20 dominant genera in the rhizosphere of control and inoculated plants at 13 dpi is shown in Fig. 2c. The results show that the pathogen largely dominated the rhizosphere microbiota. Relative abundance of *Ralstonia* genus significantly increased in inoculated plants from both the resistant and susceptible genotypes (6.01% and 16.41%, respectively) compared to control plants (1.29% and 2.04%, respectively). Furthermore, rhizosphere bacterial microbiota of control plants of the resistant and susceptible genotypes exhibited significant differences in the relative abundance of the following dominant bacterial genera: *Dyella* (4.02 and 0.88%, respectively), *Rhodococcus* (3.81 and 1.03%, respectively), *Burkholderia-Caballeronia-Paraburkholderia* (3.7 and 1.61%, respectively), *Sphingobacterium* (2.15 and 0.0%, respectively), *Chryseobacterium* (2.02 and 0.0%, respectively), *Pseudomonas* (1.4 and 5.28%, respectively), *Siccibacter* (1.15 and 2.57%, respectively) (Table S1 and Table S2).

### 3.3. Comparison of rhizosphere bacterial communities across *S. malmeanum* accessions

Unique and shared ASVs were associated to each treatment or genotype based on their presence in at least 2 of the 3 biological replicates (Fig. 3a). A total of 403 ASVs were shared across all conditions, indicating the presence of a core rhizosphere bacterial community that persisted regardless of genotype or pathogen infection. Control plants showed a higher number of unique ASVs compared to inoculated plants, and the resistant genotype exhibited the highest number of unique ASVs (309).

**Fig 3.**
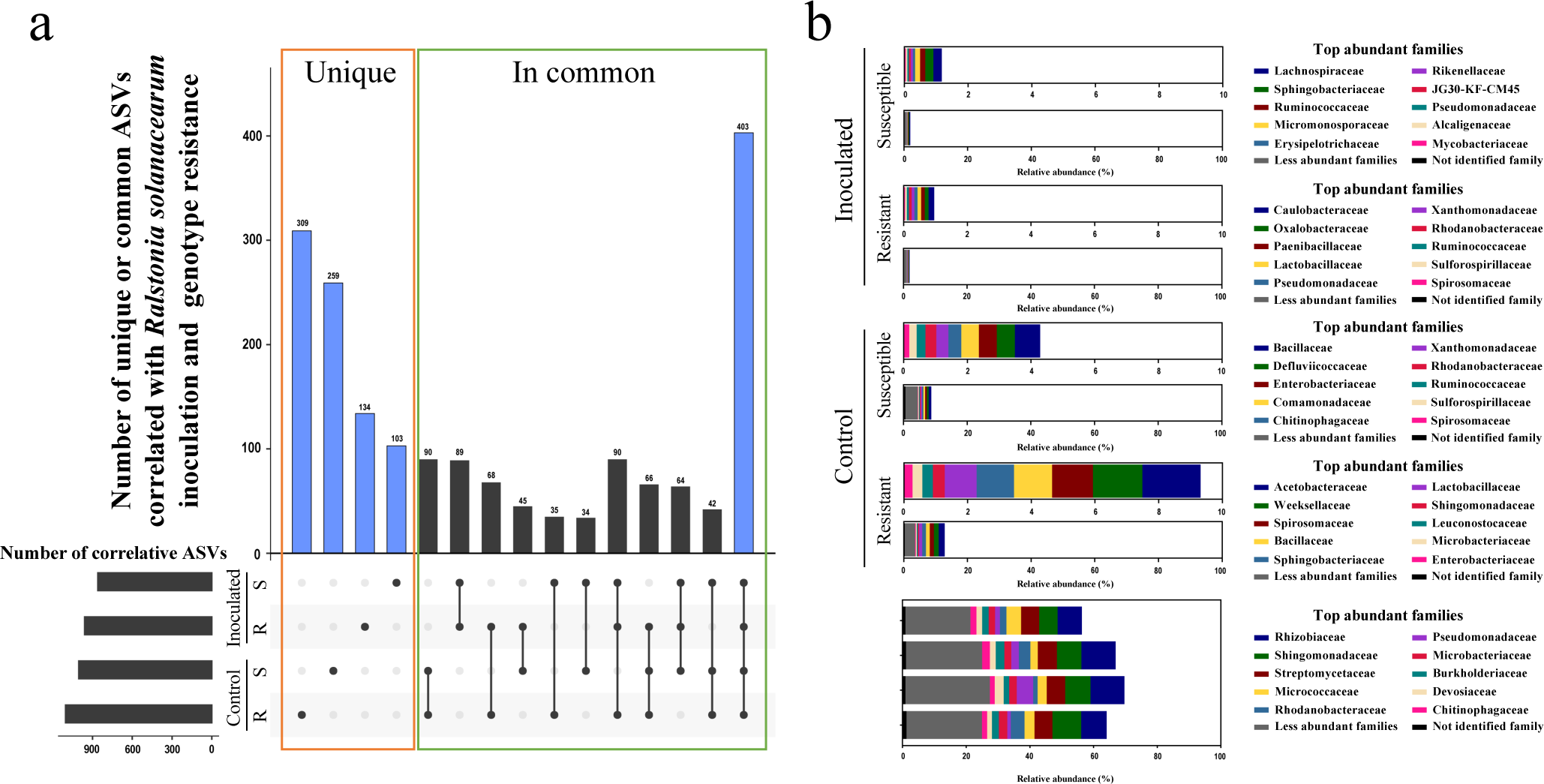
a. UpSet plot showing taxa significantly correlated with *Ralstonia solanacearum* inoculation, with control conditions and genotype resistance 13 days post inoculation. The numbers on the top of columns indicate the number of unique or common correlated ASVs among different genotypes and treatments. Blue columns to the left represent unique ASVs correlated with different samples. The blue column to the right represents common correlated ASVs among all samples. The horizontal bars to the left indicate the number of ASVs correlated with control or inoculated samples from the resistant (R) or the susceptible (S) genotype. The dots and lines below the diagram indicate the intersecting relationship of samples shown in the columns above. b. Relative abundance (%) of the major bacterial families in the rhizosphere microbiota of samples related with unique ASVs correlated with different samples or common correlated ASVs among all samples (blue columns in a).

To further characterize taxonomic patterns underlying shared and genotype-specific ASVs, the relative abundance of the major bacterial families in the rhizosphere microbiota was analyzed (Fig. 3b). The ten most abundant families shared across all accessions and treatments were Rhizobiaceae (36.88%), Sphingomonadaceae (26.41%), Streptomycetaceae (23.07%), Micrococcaceae (12.86%), Rhodanobacteraceae (11.37%), Pseudomonadaceae (10.09%), Microbacteriaceae (9.26%), Burkholderiaceae (8.75%), Devosiaceae (7.93%) and Chitinophagaceae (7.67%) (Table S3).

In control plants of the resistant genotype, a distinct set of families contributed to the genotype-specific fraction of the rhizosphere microbiota. The ten most abundant families were Acetobacteraceae (1.83%), Weeksellaceae (1.56%), Spirosomaceae (1.28%), Bacillaceae (1.18%), Sphingobacteriaceae (1.18%), Lactobacillaceae (0.99%), Sphingomonadaceae (0.37%), Leuconostocaceae (0.33%), Microbacteriaceae (0.31%) and Enterobacteriaceae (0.28%) (Fig. 3b and Table S4). Following pathogen inoculation, only two of these families, Lactobacillaceae (0.11%) and Spirosomaceae (0.01%), remained within the ten most abundant families in the rhizosphere of the resistant genotype. The remaining families were no longer detected after inoculation or decreased markedly in relative abundance, shifting into the less abundant family group. In the susceptible genotype, control plants displayed a different set of genotype-specific dominant families, including Bacillaceae (0.79%), Defluviicoccaceae (0.57%), Enterobacteriaceae (0.57%), Comamonadaceae (0.56%), Chitinophagaceae (0.40%), Xanthomonadaceae (0.37%), Rhodanobacteraceae (0.34%), Runimococcaceae (0.27%), Sulfurospirillaceae (0.23%) and Spirosomaceae (0.18%) (Fig. 3b and Table S5). After inoculation, only one family (Runimococcaceae, 0.11%) remained within the ten most abundant families in the rhizosphere. Bacillaceae, Defluviicoccaceae, Enterobacteriaceae, Sulfurospirillaceae and Spirosomaceae families were no longer detected, while the remaining families showed reduced relative abundance compared to control plants, switching to the less abundant families group.

### 3.4. Rhizosphere bacterial community reveals candidate taxa associated with the bacterial wilt resistant plant phenotype

Given the reduced disease progression observed in the resistant accession (Fig. 1a), further studies were prompted to identify bacterial taxa associated with resistance. Differential genera between control plants of the resistant and susceptible genotype were identified across the samples (Fig 4). The genera significantly enriched in the resistant genotype included *Cryseobacterium* (2.02%, *p* < 0.001), *Sphingobacterium* (2.15%, *p* < 0.001)*, Komagataeibacter* (0.85%, *p* < 0.001)*, Gluconobacter* (0.81%, *p* < 0.001)*, Lactobacillus* (1.37%, *p* < 0.001)*, Dyella* (4.02%, *p* < 0.001), *Burkholderia-Caballeronia-Paraburkholderia* (3.70%, *p* = 0.004), *Dyadobacter* (1.70%, *p* < 0.006) and *Rhodococcus* (3.81%, *p* = 0.021) (Table S2). Several of these genera (*Burkholderia-Caballeronia-Paraburkholderia, Dyella, Rhodococcus*, *Cryseobacterium* and *Sphingobacterium*) were also among the most abundant taxa in the rhizosphere community of resistant plants (Fig. 2c) and were also differentially abundant compared to the susceptible accession. Furthermore, specific genera of the resistant genotype were significantly different between control and inoculated plants, *Komagataeibacter* (0.85% and 0.0%, respectively)*, Gluconobacter* (0.81% and 0.0%, respectively)*, Lactobacillus* (1.37% and 0.37%, respectively), *Dyadobacter* (1.70% and 0.38%, respectively) and *Rhodococcus* (3.81% and 0.59%, respectively) (Table S6).

**Fig 4.**
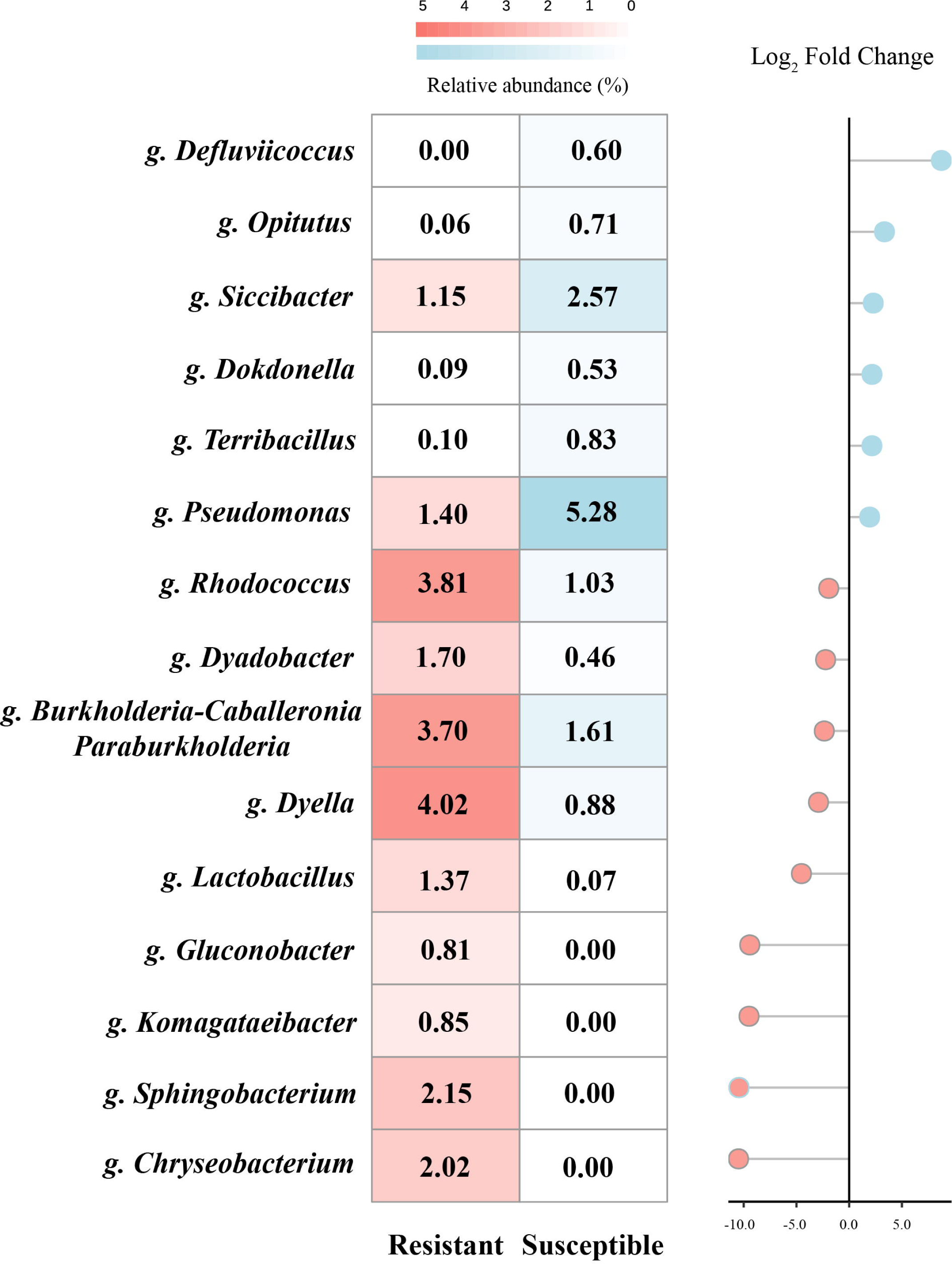
Comparison of genera between control plants of the resistant and susceptible genotype. Differentially more abundant genera found in the resistant genotype were shaded in red, while more abundant genera in the susceptible genotype were shaded in blue, with > 0.5% of relative abundance using a Zero-Inflated Gaussian Mixture Model (p < 0.05) 13 days post inoculation. The minus log_2_ fold change indicates differential genera within the resistant accession while plus log_2_ fold change indicates differential genera within the susceptible accession.

### 3.5. Bacterial functional annotation and distribution among accessions and treatments

To explore potential functional differences associated with genotype and *R. solanacearum* infection, bacterial community functions were inferred using FAPROTAX based on the classification annotations of 16S rRNA sequences. This analysis identified 50 functional groups, encompassing 225 genera, which represent 30.8% of all assigned genera (Fig. 5 and Table S7).

**Fig 5.**
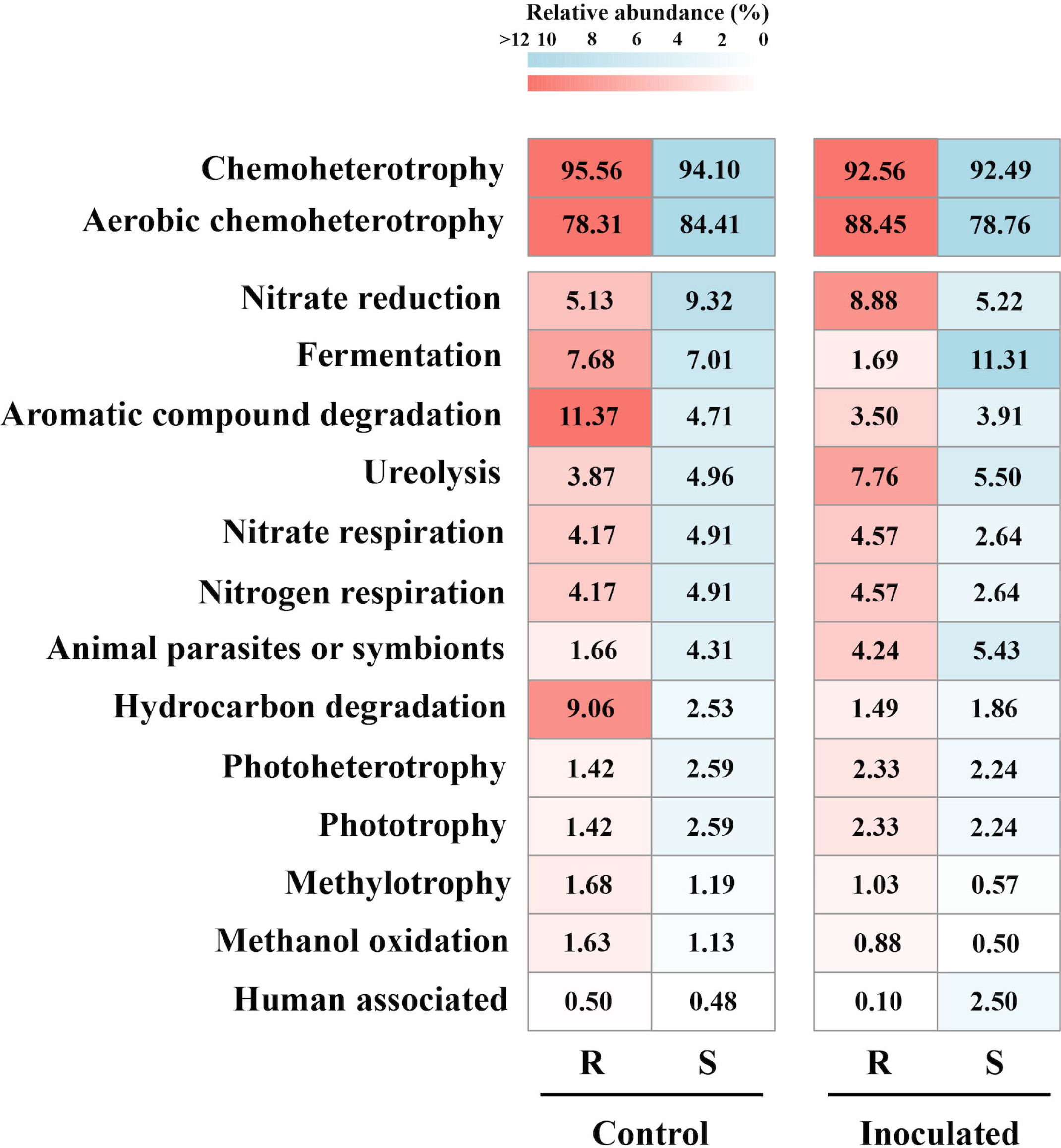
Heatmap showing FAPROTAX functional assignment of the rhizosphere bacterial community. ASVs occurrence (number of ASVs capable of each function) is indicated as relative abundance percentage and shaded in red for the resistant genotype and in blue for the susceptible genotype, derived from the rhizosphere of control and inoculated plants 13 days post inoculation.

Across all samples, the predominant functional groups were chemoheterotrophy and aerobic chemoheterotrophy, with average relative abundance of 93.68% and 82.48%, respectively. Comparison of inferred functional profiles revealed genotype and infection dependent patterns. Control plants of the resistant accession were enriched with hydrocarbon and aromatic compound degradation functional groups. Followings pathogen inoculation, these functional groups decreased along with fermentation functional groups. In contrast, the susceptible genotype showed a relative enrichement of fermentation functional groups, whereas functions related to nitrogen and nitrate respiration decreased.

### 3.6. Co-occurrence network analysis reveals putative keystone taxa in the *Solanum malmeanum* rhizosphere

To explore microbial interactions potentially associated with plant genotype, co-occurrence networks were generated for resistant and susceptible genotypes under control and inoculated conditions. Both networks exhibited marked differences in structure and complexity (Fig. 6). The network associated with the resistant genotype was composed of 62 nodes and 55 edges, whereas the susceptible genotype network contained 129 nodes and 598 edges, indicating a higher level of microbial connectivity and network density in the susceptible genotype. Additionally, the resistant genotype network exhibited 44 positive and 11 negative associations, compared with 384 positive and 214 negative associations in the susceptible genotype. The modularity was higher in the resistant network (0.87) compared to the susceptible (0.42), suggesting a more compartmentalized structure (Fig. 6a and Table S8). The co-occurrence analysis revealed both positive and negative interactions between *R. solanacearum* and specific bacterial taxa, with distinct patterns observed between the resistant and susceptible genotypes. In the resistant genotype, *R. solanacearum* showed a positive association with the genus *Novosphingobium*, whereas negative associations were detected with genera *Azospirillum*, *Acidovorax*, and members of the family Sphingomonadaceae. In contrast, in the susceptible genotype, *R. solanacearum* was positively associated with *Microbacterium*, *Bacillus*, and members of the family Enterobacteriaceae. Meanwhile, negative associations were identified with *Burkholderia*, *Ochrobactrum*, *Pseudomonas*, *Azospirillum* and the family Enterobacteriaceae, reflecting a more complex and heterogeneous interaction landscape.

**Fig 6.**
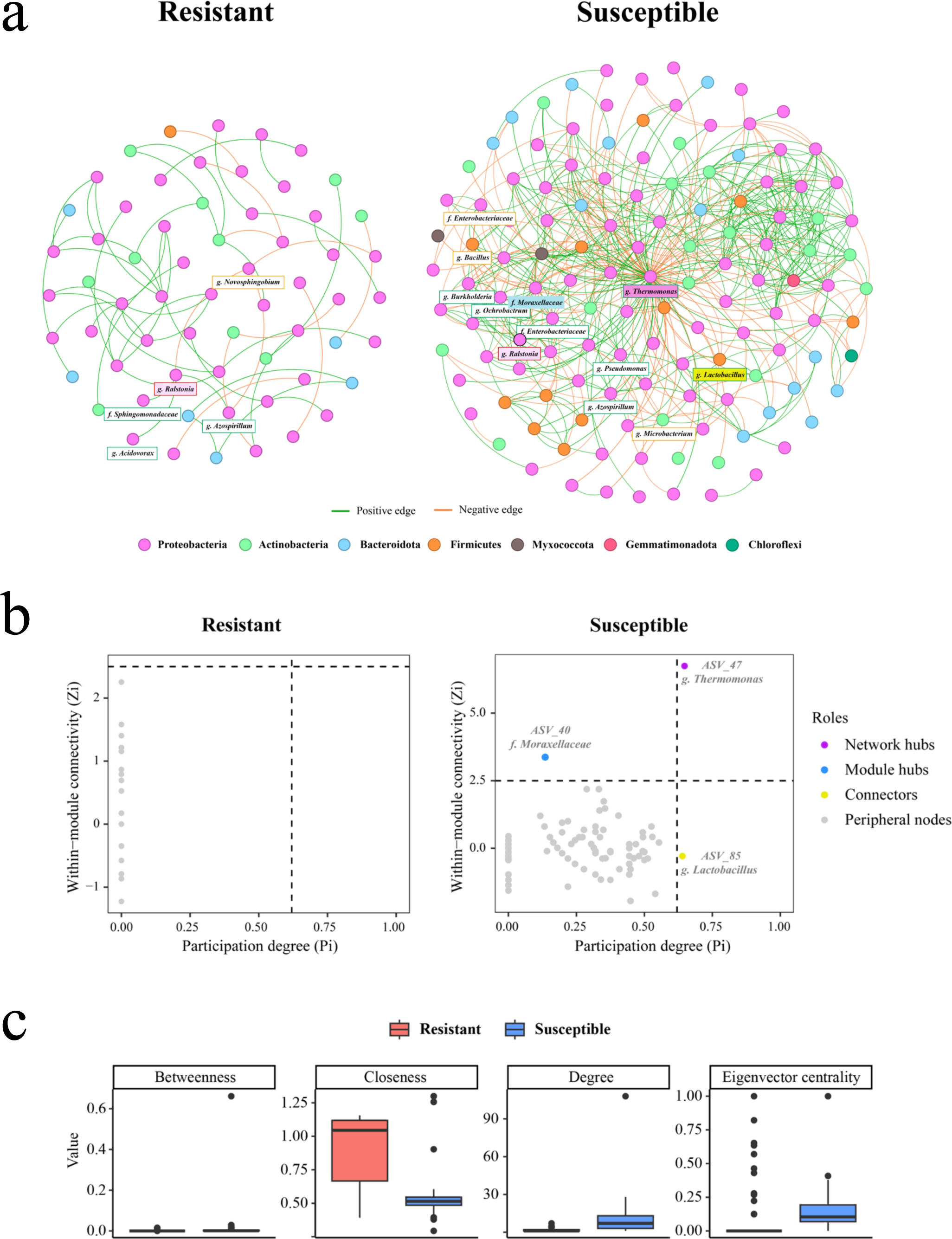
a. Co-occurrence network of rhizosphere bacterial community in genotypes of wild potato *Solanum malmeanum* resistant and susceptible to *Ralstonia solanacearum.* Each node represents different amplicon sequence variants (ASVs), while the edges (lines) indicate correlation between the nodes. Edges between nodes are colored green for positive correlations and orange for negative correlations. The names of bacterial taxa with positive and negative interactions with *Ralstonia solanacearum* (highlighted in red) were framed in green and orange respectively. The network hub, module hub and connector were indicated with purple, light blue and yellow frames, respectively. b. Topological roles within the bacterial networks. Each dot represents a node and threshold values of Zi and Pi for classifying nodes were 2.5 and 0.62 respectively, as defined by Olesen et al. (2007). c. Network parameters (betweenness, degree, closeness and Eigenvector centrality) for the networks of the resistant (Red) and susceptible (Blue) genotypes.

Topological role analysis within the co-occurrence networks revealed notable differences in the presence of keystone taxa between genotypes (Fig. 6b). In the resistant genotype, no network or module hubs were identified, and all ASVs were classified as peripheral nodes. In contrast, the susceptible genotype network exhibited a more complex architecture with the presence of topological hubs. The genus *Thermomonas* was identified as a network hub, indicating a central role in connecting multiple modules and potentially influencing global network structure. Additionally, a module hub was identified in the family Moraxellaceae, suggesting that this taxon may coordinate interactions within a subcommunity. Finally, the genus *Lactobacillus* was classified as a connector, linking multiple modules without being central within any of them. The presence of these key nodes in the susceptible network may indicate a more modular and hierarchical structure. Global network metrics further supported structural differences between genotypes. The average node degree was significantly lower in the resistant genotype (1.77) than in the susceptible genotype (9.27) (*p* < 0.001). Similarly, closeness was significantly higher in the resistant network (0.891) compared to the susceptible one (0.535) (*p* < 0.001). In contrast, betweenness did not differ significantly between networks (*P* = 0.15), however, it was nearly an order of magnitude lower in the resistant genotype, suggesting a trend toward reduced dependence on intermediary nodes (Fig. 6c and Table S9).

## 4. Discussion

Plant genotypes are known to recruit distinct microbial communities, but it remains unclear whether tolerance to soilborne pathogens depends mainly on intrinsic plant traits or on the protective capacity of the microbiota they assemble. In this study we investigated whether resistance of the wild potato *S. malmeanum* to bacterial wilt is associated with differences in the composition, structure, and functional potential of the rhizosphere bacterial community. The rhizosphere microbiota is complex and largely dependent on the surrounding soil environment. Its structure is shaped by a range of biotic and abiotic factors, with soil type and physicochemical properties being the primary determinants (Berg and Smalla, 2009; İnceoğlu et al., 2011). The findings of this study provide new insights into the dynamics of *R. solanacearum* colonization and the rhizosphere bacterial communities of wild potato accessions with different resistance levels. Interestingly, despite lower pathogen concentrations in the resistant genotype at 13 dpi, pathogen levels were comparable between genotypes by 27 dpi. This temporal pattern suggests that resistance in *S. malmeanum* may not rely on complete exclusion of the pathogen from the rhizosphere, but rather on delayed colonization or reduced disease progression. Such dynamics underscore the complexity of plant-pathogen interactions, where pathogen suppression may be transient and modulated by environmental or host-related factors (Korenblum et al., 2020). Correlations have been previously identified, in which higher bacterial diversity was associated with an increased suppression of *R. solanacearum*, emphasizing the role of rhizosphere microbial communities in disease regulation Correlations have been previously identified, in which higher bacterial diversity was associated with an increased suppression of *R. solanacearum* (Hu et al., 2016; Wang et al., 2017; Deng et al., 2020).

Amplicon sequencing analysis revealed marked shifts in rhizosphere bacterial community composition in response to pathogen inoculation and plant genotype. The most important differences were observed at 13 dpi between inoculated and control plants, and between resistant and susceptible genotypes at both sampling times. The dominance of *Ralstonia* in inoculated treatments highlights its strong competitive ability to colonize the rhizosphere, potentially displacing other microbial taxa. This finding is consistent with previous reports indicating that pathogen invasion imposes selective pressures on rhizosphere microbiota, altering community stability and composition (Berendsen et al., 2012; Pascale et al., 2020). Despite these perturbations, resistant and susceptible *S. malmeanum* accessions maintained distinct rhizosphere bacterial community compositions. The resistant genotype remained asymptomatic throughout the experiment, and beta diversity analysis also suggest that the rhizosphere microbiota associated with the resistant genotype may contribute to disease modulation. Notably, it contained unique bacterial genera that were already enriched under control conditions, including *Dyella*, *Rhodococcus*, *Burkholderia-Caballeronia-Paraburkholderia*, *Chryseobacterium* and *Sphingobacterium*. These genera have been previously associated with plant growth promotion, pathogen suppression, and soil health, supporting their potential role in modulating resistance to bacterial wilt (Compant et al., 2005; Mendes et al., 2011; Belov et al., 2018; Hakim et al., 2021; Jung et al., 2023).

Analysis of unique and common ASVs, further indicated that control treatments showed a higher number of unique taxa than inoculated treatments, particularly in the resistant genotype. This suggests that inoculation with *R. solanacearum* produces a selective pressure that reduces rhizosphere microbial diversity, consistent with previous observations in tomato, where pathogen invasion disrupted rhizosphere bacterial communities leading to a reduction in the diversity and abundance of non-pathogenic bacteria (Wei et al., 2018). The higher number of unique ASVs in the resistant genotype under control conditions may reflect a more diverse or specialized microbial reservoir capable of contributing to disease suppression.

Differential functions are displayed by unique families identified among the top ten most abundant in control plants of the susceptible and resistant genotypes. In the susceptible genotype, unique families are associated with functions related to organic matter degradation, nutrient cycling, and environmental resilience, such as Defluviicoccaceae, Comamonadaceae, Chitinophagaceae, Xanthomonadaceae, Rhodanobacteraceae, and Spirosomaceae (Khan et al., 2002; Chang and Zylstra, 2010; Cutiño-Jiménez et al., 2020; Bessarab et al., 2022; Kim et al., 2024). While these families contribute to soil functioning, their roles in direct disease suppression are less clear. In contrast, control plants of the resistant genotype were enriched in families such as Bacillaceae, Sphingobacteriaceae, and Acetobacteraceae, which are frequently with biocontrol activities and disease suppression. Their roles often involve the production of antimicrobial compounds or competitive exclusion of pathogenic microbes (Saravanan et al., 2008; Shafi et al., 2017; Ajesh et al., 2024). Additionally, families such as Lactobacillaceae and Enterobacteriaceae may contribute indirectly to plant health by enhancing overall microbial community balance or promoting plant growth (Jha et al., 2011; Jaffar et al., 2023). Enterobacteriaceae has been associated with resistance against *R. solanacearum* in tomato (Roy et al., 2019).

Two of the differential genera enriched in the resistant genotype, *Komagataeibacter* and *Gluconobacter,* belong to Acetobacteraceae, an acetic acid bacterial family. Members of this family are known for acetic acid fermentation and tolerance to harsh environmental conditions, and *Gluconobacter* species have been reported to inhibit fruit and plant pathogens (Rashad and Moussa, 2020; Delgado et al., 2021). Studies have shown that *Gluconobacter* exhibits inhibitory effects against fruit and plant pathogens, demonstrating its potential utility in disease management (Rashad and Moussa, 2020; Delgado et al., 2021). On the other hand, *Komagataeibacter* species are distinguished by their role in acetic acid fermentation and their resilience to environmental stressors as low pH, high ethanol concentrations, and high osmolarity (Nascimento et al., 2021). Although, direct antagonistic effects against *R. solanacearum* have not been demonstrated for these genera, protective effects have been reported for related taxa such as *Gluconacetobacter diazotrophicus* (Rodriguez et al., 2019; Srebot et al., 2024). Similarly, studies have revealed that *Chryseobacterium,* another differential genus found in the resistant genotype in this work, effectively reduces bacterial wilt; in tomato (Huang et al., 2017; Yoo et al., 2020). Moreover, some species within this genus and *Burkholderia-Caballeronia-Paraburkholderia* have been recognized for their capacity to degrade aromatic compounds, a functional group that was enriched in the resistant genotype (Goris et al., 2004; Jung et al., 2023). Additionally, *Sphingobacterium* species are notable for their potential to survive in extreme environments and their ability to suppress fungal pathogens (Xu et al., 2020; Ajesh et al., 2024). *Lactobacillus* species, known for their probiotic properties, have been explored for their potential in biocontrol applications, particularly against various bacterial plant pathogens. Specific studies on *Lactobacillus* strains targeting *R. solanacearum*, are limited to tomato plants and focused on the role of defense enzymes and antibacterial compounds in conferring disease resistance (Konappa et al., 2016; Murthy et al., 2016). Certain *Rhodococcus* species have demonstrated potential in managing plant diseases, by disrupting the quorum sensing system of bacteria. This genus is well-known for its ability to degrade a wide range of hydrocarbons, including alkanes, aromatic compounds, and other organic pollutants (Barbey et al., 2013). The enrichment of these taxa is consistent with the observed increase in functional groups related to hydrocarbon and aromatic compound degradation in the resistant genotype, suggesting a functional link between microbial metabolism and disease suppression.

Functional predictions revealed additional genotype-dependent responses to pathogen inoculation. In the resistant genotype, nitrate reduction functions increased after inoculation, whereas these functions declined in the susceptible genotype. Microbial nitrate nutrition can influence nitrogen availability and signaling in the rhizosphere, potentially affecting plant health and disease resistance. However, the specific impacts of microbial nitrate reduction on plant-pathogen interactions are complex and can vary depending on the plant species, pathogen, and environmental conditions (Fu et al., 2018; Liu et al., 2023).

Among the differential genera identified in the resistant genotype, some are associated with biocontrol activities. However, only a few have demonstrated antagonistic effects against bacterial wilt, with the limited evidence primarily derived from studies in tomato (Konappa et al., 2016; Murthy et al., 2016; Huang et al., 2017; Roy et al., 2019; Yoo et al., 2020). The identification of specific bacterial families and genera linked to disease suppression in the resistant genotype highlights potential candidates for microbial inoculants or bioindicators of soil health. A more detailed characterization of the functional roles of these taxa in pathogen resistance within the rhizosphere microbiome, will be necessary to clarify the mechanisms underlaying these associations. In this context, complementary approaches such as metagenomic or transcriptomic analyses could provide a deeper understanding of the dynamic interplay between plants, pathogens, and microbiota.

Beyond differences in taxonomic composition and predicted functions, co-occurrence networks analysis revealed clear structural and functional differences in the rhizosphere microbiota associated with resistant and susceptible wild potato genotypes. The lower complexity and connectivity observed in the resistant genotype, evidenced by fewer nodes and edges, a lower average degree, and the absence of central hubs, suggests a more streamlined microbial network. Despite this apparent simplicity, this network showed higher modularity and closeness, suggesting a compartmentalized organization with efficient internal connectivity. Such organization has been proposed to enhance the functional stability and resilience of the community to pathogen invasion (Coyte et al., 2015). Importantly, the lack of keystone taxa (network or module hubs) in the resistant genotype may reflect a less centralized network, reducing dependency on dominant nodes and potentially contributing to community robustness. In contrast, the susceptible genotype exhibited a more complex and hierarchical network, which suggests specialized roles and higher interaction diversity. It may also indicate increased vulnerability to disruption, especially under pathogen pressure (Toju et al., 2018).

Interestingly, our results contrast with previous studies reporting higher network complexity in healthy or resistant plants (Xiao et al., 2024; Lai et al., 2025), our findings revealed the opposite pattern. This discrepancy may reflect host-specific or environment-dependent differences in microbiome assembly. For instance, it has been suggested that the recruitment of highly specialized microbial consortia by resistant genotypes may result in tightly filtered but functionally effective communities, even if these are structurally simpler (Toju et al., 2018; Trivedi et al., 2020).

Overall, the observed differences in community composition, functional potential, and microbial interaction networks suggest that resistance to bacterial wilt in wild potato is associated with distinct rhizosphere ecological configurations rather than the action of individual microbial taxa. These insights highlight the potential of leveraging microbial network analysis to guide the development of microbiome-informed crop protection strategies.

## 5. Conclusion

This study demonstrates that resistance to *R. solanacearum* in wild potato *S. malmeanum* is associated with distinct rhizosphere microbial communities and functional traits that are shaped by plant genotype. Notably, resistant plants harbored specific bacterial taxa and interaction patterns already under non-inoculated conditions, indicating a genotype-dependent recruitment of rhizosphere microbiota. These microbial assemblages were further associated with traits linked to disease suppression and ecological stability. Together, these findings reinforce the importance of considering the rhizosphere microbiome as an integral component of resistance phenotypes and provide a basis for exploring microbiome-based strategies for disease management.

## Acknowledgements

M.V.F. acknowledges funding from Carlos Vaz Ferreira Grant and the Basics Sciences Development Program (PEDECIBA) from Uruguay. We thank technical staff from the Biotechnology Laboratory in INIA (Uruguay) and Southern Regional Center (Universidad de la República, Uruguay) for assistance with in vitro plant propagation and plant monitoring. G.L. acknowledges the supercomputing infrastructure of Soroban (SATREPS MACH—JPM/JSA1705) at Centro de Modelación y Computación Científica at Universidad de La Frontera.

