## Supplemental Figure S1 for "Bacterial wilt resistance is correlated with rhizosphere bacterial communities in wild potato *Solanum malmeanum*"

**Supplementary Figure**

**Figure S1.**

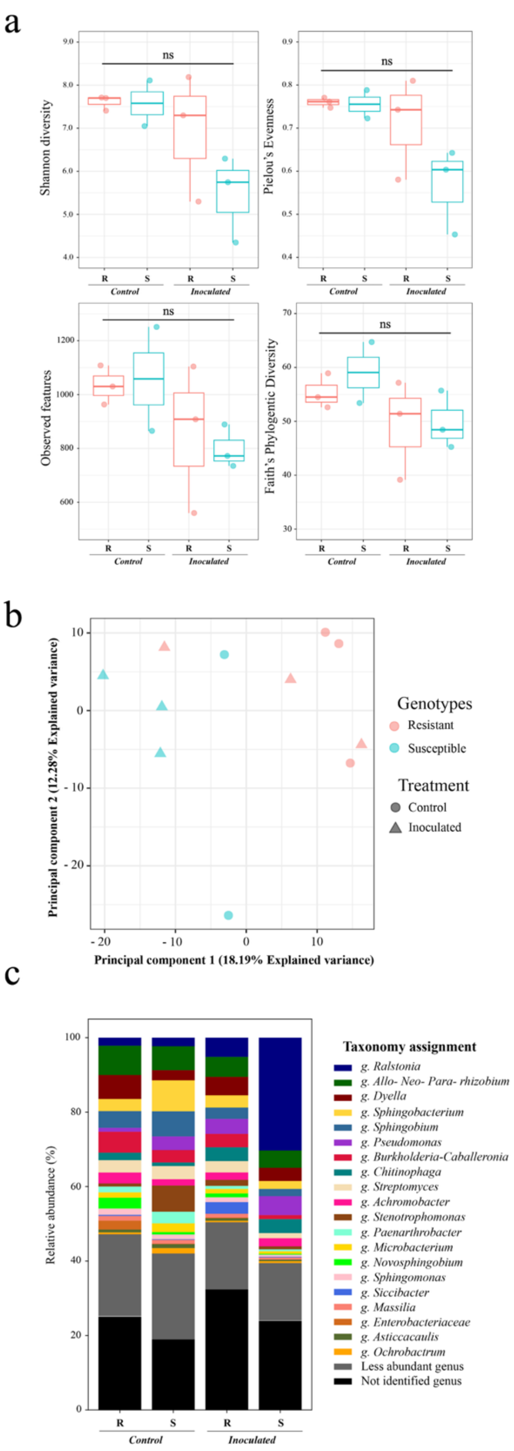


**Fig S1.** a. Comparative alpha-diversity analysis of rhizosphere bacterial community among the bacterial wilt resistant (R) and susceptible (S) genotype 27 days post inoculation.

Significance of Shannon diversity index, Faith's phylogenetic index, Pielou's Evenness index and observed features (number of ASVs) were calculated with Kruskal-Wallis test ($p < 0.05$). ns = not significant. b. Comparative beta-diversity analysis of rhizosphere bacterial community between the bacterial wilt resistant and the susceptible genotype 27 days post inoculation. The distribution pattern of control and inoculated plants from the resistant and susceptible genotype were normalized using Aitchison distance. Significance of each factor and its interaction was evaluated using the PERMANOVA test ($p < 0.05$). c. Relative abundance (%) of the major bacterial genera in the rhizosphere microbiota of the resistant (R) and susceptible (S) genotypes 27 days post inoculation.
