## Supplemental Table S9 for "Bacterial wilt resistance is correlated with rhizosphere bacterial communities in wild potato *Solanum malmeanum*"

t.test(Betweenness ~ Dataset, data_combined)
data:  Betweenness by Dataset
t = -1.4421, df = 129.4, p-value = 0.1517
alternative hypothesis: true difference in means between group resistant and group susceptible is not equal to 0
95 percent confidence interval:
  -0.017581144  0.002757024
sample estimates:
  mean in group resistant mean in group susceptible 
0.0008901816              0.0083022417 

t.test(Closeness ~ Dataset, data_combined)
data:  Closeness by Dataset
t = 10.079, df = 80.269, p-value = 6.531e-16
alternative hypothesis: true difference in means between group resistant and group susceptible is not equal to 0
95 percent confidence interval:
  0.2859677 0.4266607
sample estimates:
  mean in group resistant mean in group susceptible 
0.8908800                 0.5345658

t.test(degree ~ Dataset, data_combined)
data:  degree by Dataset
t = -7.6745, df = 134.36, p-value = 3.036e-12
alternative hypothesis: true difference in means between group resistant and group susceptible is not equal to 0
95 percent confidence interval:
  -9.429179 -5.565070
sample estimates:
  mean in group resistant mean in group susceptible 
1.774194                  9.271318
